# 3D-PAULM: Integrated Photoacoustic Tomography and Ultrasound Localization Microscopy for Multiparametric Brain and Tumor Imaging

**DOI:** 10.64898/2026.04.30.722008

**Authors:** Yirui Xu, Rui Yao, Huaxin Sheng, Nanchao Wang, Xinyuan Yu, Xueyi Cai, Jiaji Cai, Jinhuan Luo, Jingting Li, Wei Yang, Pengfei Song, Vladislav V. Verkhusha, Junjie Yao

## Abstract

Understanding processes such as blood–brain barrier (BBB) disruption and tumor progression can greatly benefit from simultaneous molecular, functional, and hemodynamic imaging in deep tissue, yet few existing imaging modalities can provide all three in a single system. Here, we present an integrated imaging platform that combines 3D photoacoustic tomography with ultrasound localization microscopy (3D-PAULM) to enable intrinsically co-registered, multiparametric imaging. 3D-PAULM unifies multispectral photoacoustic molecular imaging, ultrasound B-mode imaging, microbubble-enhanced power Doppler, and ultrasound localization microscopy, and concurrently measures blood oxygenation, blood perfusion, microvascular flow dynamics, and molecular probes from near-infrared dyes and photoswitchable phytochromes. We apply 3D-PAULM to quantify BBB leakage in focal ischemia and systemic inflammation, and to perform high-sensitivity molecular imaging of solid tumors alongside functional mapping of tumor hypoxia and super-resolved vascular remodeling. Together, these results establish 3D-PAULM as a versatile platform for integrated functional and molecular imaging in deep tissue.

## Introduction

Multimodal imaging is increasingly important in preclinical research, where understanding disease progression and therapeutic response requires concurrent anatomical, functional, and molecular information. Processes such as blood–brain barrier (BBB) disruption and tumor progression are tightly coupled at molecular and hemodynamic levels. Yet existing modalities still struggle to provide simultaneous, three-dimensional molecularly specific imaging together with complementary hemodynamic characterization in deep tissue [1–4]. Overcoming this gap would enable direct interrogation of barrier dysfunction, perfusion deficits, and treatment response within a unified experimental framework.

Photoacoustic tomography (PAT) combines optical absorption contrast with deep-tissue acoustic detection, enabling visualization of endogenous and exogenous optical absorbers at centimeter-scale depths [5–7]. Because hemoglobin is the dominant optical absorber in tissue in the visible and near-infrared wavelength range, PAT is particularly well suited for functional imaging of blood oxygen saturation (sO₂), a widely used readout of tissue oxygenation and metabolic state [8–11]. PAT is also naturally compatible with ultrasound (US) imaging because both rely on acoustic detection. In integrated PA/US systems, ultrasound provides complementary anatomy and high-temporal-resolution sensitivity to blood flow; Doppler methods capture bulk flow, and gas-filled microbubbles can further boost power Doppler (PWD) sensitivity to vascular perfusion [12, 13]. This shared detection geometry yields intrinsic PA/US co-registration and enables multiparametric imaging without post hoc alignment.

Despite these strengths, conventional PA/US imaging remains limited in two important ways for deep-tissue preclinical studies. First, spatial resolution is constrained by the acoustic diffraction limit, which restricts the ability to resolve fine vascular structure and flow dynamics. Higher-frequency transducers and other system-level optimizations can improve resolution and volumetric performance, but they typically do so at the expense of penetration depth and remain diffraction limited [14, 15]. Second, endogenous absorbers, especially hemoglobin, often dominate the photoacoustic signal and obscure weak signals from exogenous molecular probes, reducing in vivo molecular sensitivity. Approaches such as spectral unmixing and bright exogenous agents can help when probe signals are strong and spectrally distinct from background absorbers [5, 16, 17], but they are less effective for weakly absorbing probes such as genetically encoded reporters. Accordingly, there remains a need for integrated strategies that combine super-resolution hemodynamic imaging with background-suppressed molecular detection in deep tissue.

To overcome the resolution barrier, we integrated PAT with three-dimensional ultrasound localization microscopy (ULM) to create 3D-PAULM, which enables super-resolution imaging of vascular structure and hemodynamics in deep tissue. By localizing and tracking individual microbubbles, ULM surpasses the diffraction limit and resolves vascular organization as well as blood-flow speed and direction at sub-diffraction resolution [18–20]. Incorporating ULM therefore expands the hemodynamic information available from PAT to include microvascular architecture, flow dynamics, and oxygenation within a single framework. Comprehensive multiparametric imaging can be completed in under one minute per scan position.

To address molecular sensitivity, we also incorporated exogenous contrast agents that provide specificity beyond endogenous absorbers. Genetically encoded probes are especially attractive because they support cell-type targeting and longitudinal *in vivo* studies [21, 22]. In particular, differential PA imaging with reversibly photoswitchable genetically encoded probes provides an effective route to deep-tissue molecular imaging with much improved sensitivity (>1000 fold) [21–23]. By toggling the probe between two spectrally distinct absorbing states, differential PA imaging isolates the switching probe signal while suppressing non-switching background absorbers. Unlike conventional spectral unmixing, this strategy does not require strong probe absorption or clear spectral separation from hemoglobin, making it well suited for weak molecular signals in blood-rich tissues such as brain and tumors. This advantage is especially important for genetically encoded probes, whose signals are often much weaker than endogenous background *in vivo*.

We thus demonstrate the utility of 3D-PAULM in two representative applications using tailored exogenous probes. First, we use a near-infrared dye that is spectrally separable from hemoglobin, indocyanine green (ICG), to quantify BBB leakage after focal ischemia and systemic inflammation. Second, we use reversibly photoswitchable bacterial phytochrome-derived probes for tumor microenvironment imaging. By reversibly switching the probe between ON and OFF states and applying differential PA imaging, we recover tumor-associated signals that would otherwise be easily overwhelmed by surrounding background.

Overall, 3D-PAULM enables longitudinal, multiparametric interrogation of disease processes by combining deep-tissue functional and molecular imaging with super-resolution hemodynamic characterization in a unified platform.

## Results

### Integrated 3D-PAULM system for co-registered multiparametric imaging

To interrogate molecular, functional, and microvascular dynamics simultaneously in deep tissue, we established 3D-PAULM, an integrated imaging platform that integrates multispectral PAT, ultrasound B-mode imaging, microbubble-enhanced power Doppler, and ULM within a single experimental framework (**Fig. 1a**). Building on our previously reported architecture [20], the present implementation incorporates interleaved PA/US imaging sequences and streamlined data transfer to support high-throughput *in vivo* studies. The interleaved acquisition scheme provides intrinsic spatial and temporal co-registration across modalities without post hoc alignment. The system operates at a PA frame rate of 30 Hz and a volumetric US frame rate of 403 Hz (**Fig. 1b**). An optimized data-streaming pipeline minimizes acquisition dead time and enables continuous streaming between frame batches, thereby avoiding temporal gaps and supporting consistent microbubble tracking for ULM. As a result, comprehensive multiparametric acquisitions, including multispectral PA and ULM, can be completed efficiently with reduced motion-induced artifacts.

**Fig. 1.**
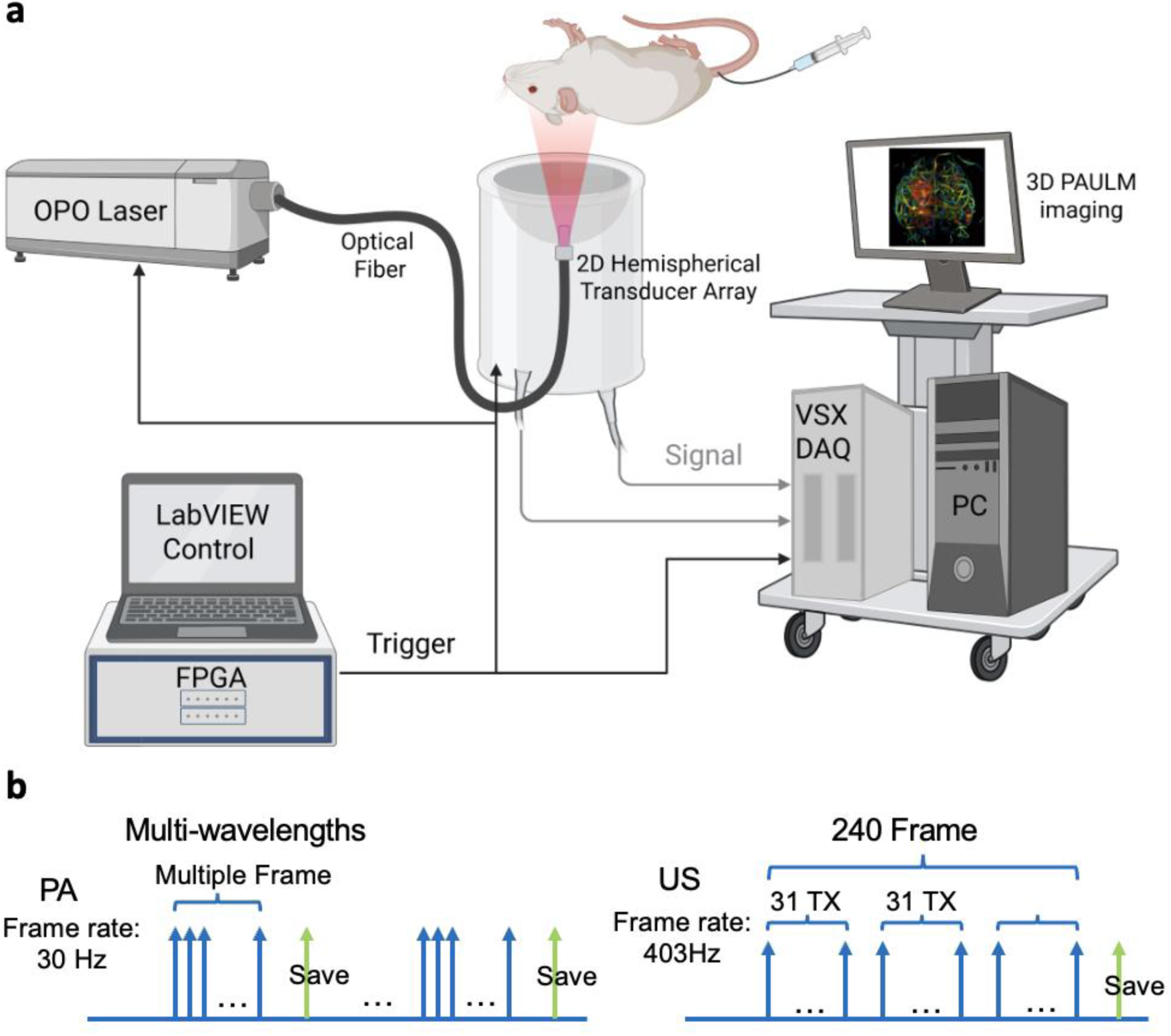
3D integrated photoacoustic tomography and ultrasound localization microscopy (3D-PAULM). **(a)** Schematic of the imaging system setup. The 3D-PAULM system comprises a data acquisition system, a short-pulsed OPO laser, and a customized 2D spherical ultrasound transducer array. The fiber outlet is positioned at the center of the array for photoacoustic (PA) illumination. Laser firing, ultrasound detection, and data acquisition are synchronized by a LabVIEW-based FPGA module. During imaging, the transducer array is immersed in water. The imaging target is positioned around the focal zone of the transducer array (approximately 40 mm from the transducer surface) and raster scanned using a 2D motorized translation stage. The system can perform multispectral PA imaging, ultrasound (US) B-mode imaging, microbubble-enhanced US power Doppler imaging (PD), and ultrasound localization microscopy (ULM). **(b)** Data acquisition sequence of PA and US imaging with respective volumetric frame rates of 30 Hz and 403 Hz.

Together, these features enable simultaneous quantification of hemoglobin oxygenation, blood perfusion, microvascular flow dynamics, and molecular contrast from both exogenous dyes and genetically encoded probes. This framework enables the *in vivo* studies presented below, including integrated characterization of BBB disruption and tumor microenvironments.

### Multispectral PA imaging enables volumetric quantification of BBB disruption

To assess BBB leakage *in vivo*, we used indocyanine green (ICG) as an NIR contrast agent in multispectral PA imaging (**Fig. 2a**). ICG is a clinically approved dye that binds plasma proteins and, when the BBB is intact, remains intravascular and is rapidly cleared from the circulation. When the BBB is disrupted, however, ICG extravasates and clears more slowly [24, 25]. This behavior has been widely exploited in fluorescence imaging and NIR spectroscopy to probe BBB permeability under pathological conditions [26]. Here, we extend that concept to PAT by isolating ICG from surrounding blood signals and using dye extravasation as a quantitative readout of BBB leakage.

**Fig. 2.**
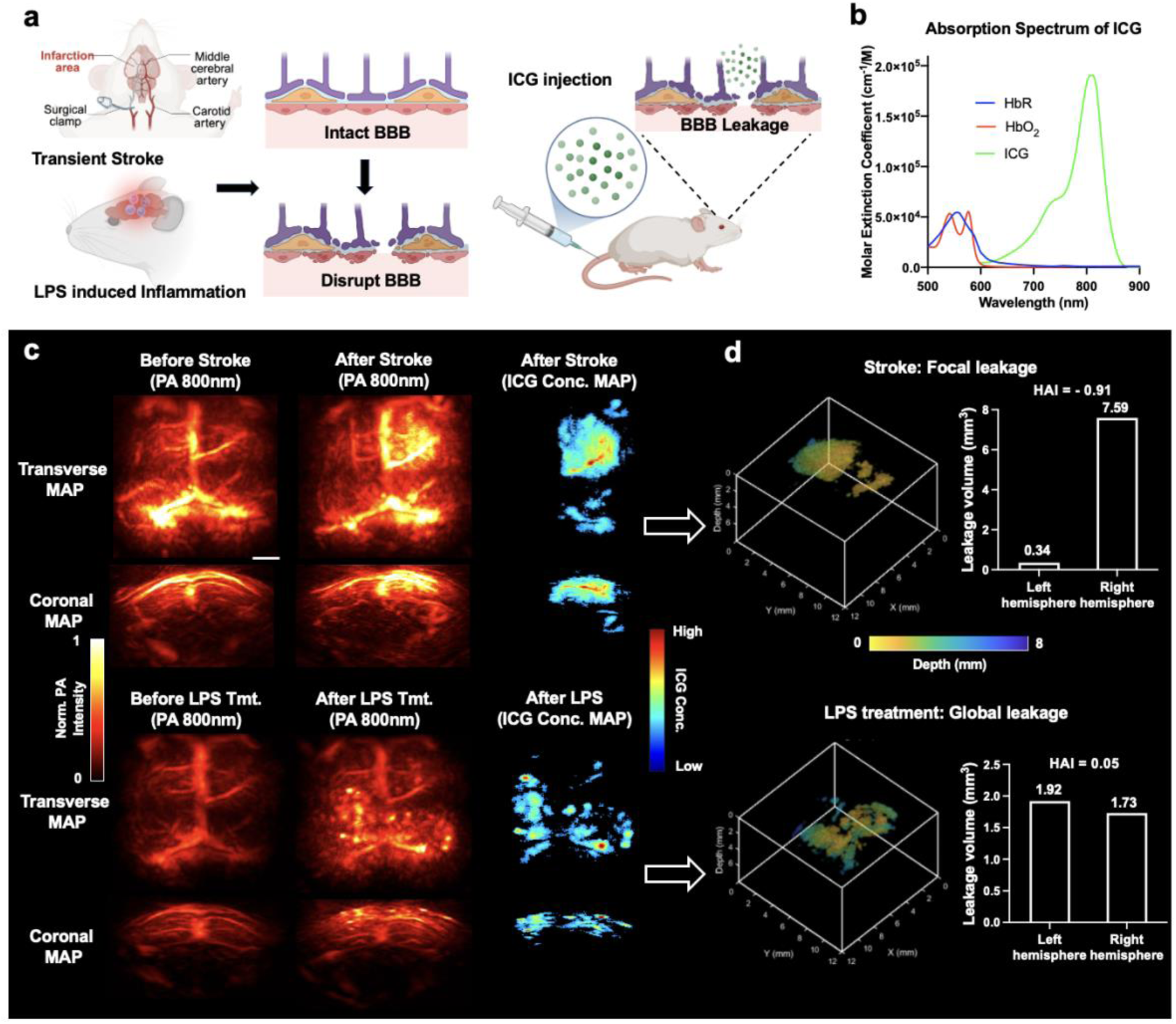
Multispectral PA imaging for BBB leakage detection. **(a)** Schematic of the experimental paradigm and BBB leakage models. BBB disruption was induced either focally by transient middle cerebral artery occlusion (stroke model) or globally by LPS-induced inflammation. After intravenous ICG injection, extravasated dye accumulates at leakage sites and becomes detectable by PA imaging. **(b)** Absorption spectra of ICG and hemoglobin (HbR/HbO₂). ICG exhibits a strong NIR absorption band that is spectrally distinct from hemoglobin. **(c)** Representative maximum-amplitude-projection (MAP) PA images before and after BBB disruption and the corresponding ICG concentration maps. Left and middle columns: single-wavelength PA images at 800 nm acquired 20 min after ICG injection before and after transient stroke (top row) or LPS treatment (bottom row). Right column: ICG concentration maps reconstructed from fluence-compensated, multispectral acquisitions (730, 800, and 870 nm) and spectral unmixing; pixels containing 0% ICG are rendered in black. Transverse (dorsal) and coronal views are shown. Scale bars, 2 mm. **(d)** Quantitative 3D analysis of ICG leakage. Top: three-dimensional rendering of ICG leakage in the stroke model, demonstrating spatially confined and hemispherically asymmetric BBB disruption. Leakage volumes were quantified separately in the left and right cerebral hemispheres, and the hemispheric asymmetry index (HAI) is reported. Bottom: three-dimensional rendering of ICG leakage after LPS treatment, revealing diffuse and bilaterally distributed BBB leakage across the brain. Corresponding leakage volumes in the left and right hemispheres are shown, together with the HAI value. Depth information is color-coded as indicated.

ICG is an effective exogenous PA contrast agent because its strong NIR absorption peak near 800 nm is spectrally well separated from hemoglobin (HbR/HbO₂) absorption (**Fig. 2b**) [27]. Accordingly, single-wavelength PA imaging at 800 nm was highly sensitive to extravascular ICG. In baseline images acquired 20 min after injection, when most circulating dye had cleared, no extravascular ICG accumulation was detected in animals with an intact BBB. In contrast, ischemic stroke or LPS treatment produced marked PA signal enhancement, consistent with ICG extravasation at sites of BBB disruption (**Fig. 2c**). To improve specificity and enable quantification, we then performed multispectral PA imaging with spectral unmixing to estimate ICG concentration. This step further separated ICG from endogenous hemoglobin background and improved localization of BBB leakage (**Fig. 2c–d**).

In the unilateral transient focal ischemia model, unmixed ICG concentration maps revealed a spatially confined leakage pattern localized mainly to the right hemisphere, consistent with the site of ischemic injury. Quantitative three-dimensional analysis identified a total leakage volume of 7.93 mm³ with pronounced hemispheric asymmetry (**Fig. 2d**, top). Leakage was concentrated overwhelmingly in one hemisphere, yielding a hemispheric asymmetry index (HAI) of −0.91, defined as HAI = (V_left_ – V_right_) / (V_left_ + V_right_), where V_left_ and V_right_ denote leakage volumes in the left and right hemispheres, respectively [28, 29]. By contrast, systemic LPS treatment produced diffuse, bilaterally distributed BBB leakage. The total leakage volume was 3.65 mm³, with similar contributions from both hemispheres and an HAI of 0.05 (**Fig. 2d**, bottom). These PA findings were consistent with post-imaging fluorescence measurements, which showed matching spatial patterns of ICG accumulation in both the stroke and LPS models (**Fig. 3a, e**, insets). Together, these results show that multispectral PA imaging enables volumetric quantitative mapping of BBB leakage and can distinguish focal disruption from diffuse, global permeability changes *in vivo*.

**Fig. 3.**
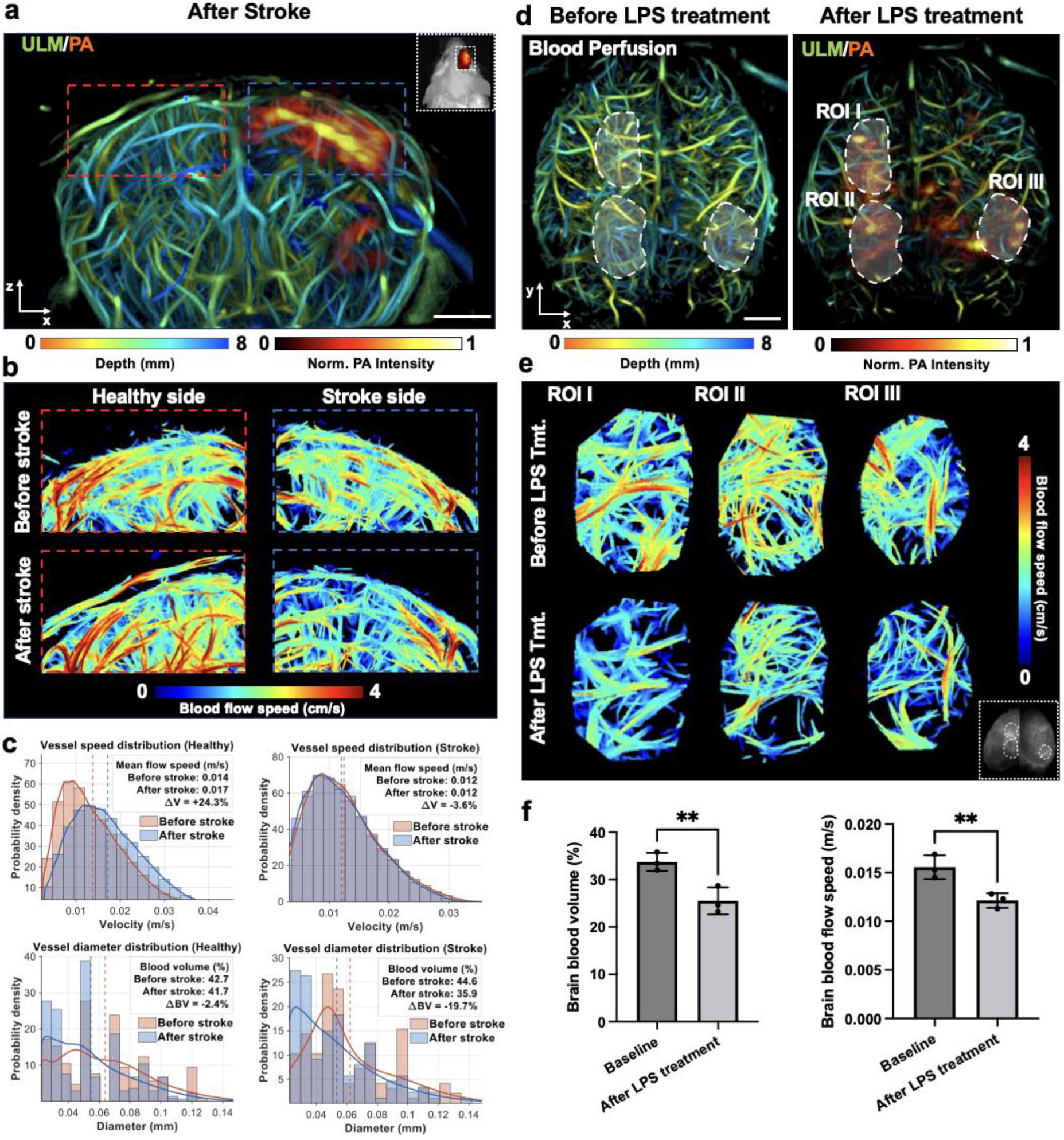
3D-PAULM of BBB leakage and associated microvascular hemodynamics in the deep brain. **(a)** Representative co-registered 3D-PAULM image acquired after transient stroke, showing super-resolved cortical microvasculature resolved by ULM and color-coded by depth, overlaid with the PA-derived ICG signal. The red and blue dashed boxes indicate representative ROIs on the healthy and stroke sides, respectively. Inset, IVIS imaging of the ICG channel. **(b)** Enlarged ULM flow-speed maps of the matched ROIs from the healthy side and stroke side, shown before and after stroke. Blood-flow speed is color-coded as indicated. **(c)** Quantitative analysis of microvascular hemodynamics within matched ROIs on the healthy and stroke sides. Top row, histograms with overlaid kernel density curves showing vessel flow-speed distributions before and after stroke. Bottom row, histograms with overlaid kernel density curves showing the corresponding estimated vessel diameter distributions. Mean flow speed and blood volume are indicated for each condition, together with relative changes between the pre-stroke and post-stroke states. **(d)** Representative baseline ULM image (left) and co-registered ULM/PA image after LPS treatment. Three representative ROIs (I–III) are outlined by dashed contours. **(e)** ULM flow-speed maps from ROIs I–III at baseline and after LPS treatment. Inset, fluorescence imaging of the ICG signal. **(f)** Statistical analysis of whole-brain blood perfusion and blood-flow speed at baseline and after LPS treatment (p < 0.05). Data are presented as mean values ± SD (n = 3 animals). A paired t-test was used.

### ULM characterization of hemodynamic change in BBB leakage region

We next used 3D-PAULM to place BBB leakage in its microvascular context by measuring vascular structure and flow dynamics together with permeability in the same co-registered volume (**Fig. 3**). In the transient stroke model, co-registered ULM and PA imaging showed that regions of BBB leakage, identified by extravascular ICG, spatially overlapped with super-resolved cerebral vasculature (**Fig. 3a**). This correspondence enabled direct comparison of leakage patterns with local microvascular features. Magnified views of matched regions of interest (ROIs) in the contralateral (healthy) and ipsilateral (stroke) hemispheres revealed pronounced microvascular reorganization after stroke (**Fig. 3b**). Quantitative analysis showed that vessel-diameter distributions in both hemispheres shifted after stroke (**Fig. 3c**), indicating redistribution toward smaller vessel calibers. This pattern is consistent with acute microvascular remodeling extending beyond the infarct core [30, 31].

Although both hemispheres exhibited structural remodeling, their hemodynamic responses differed. On the healthy side, the shift toward smaller vessels was accompanied by largely preserved blood volume (−2.4%) and increased mean flow speed (+24.3%), suggesting compensatory flow redistribution within the microvascular network [32]. On the stroke side, the same leftward diameter shift coincided with a marked reduction in blood volume (−19.7%), whereas flow-speed distributions largely overlapped before and after stroke, producing only a small change in mean flow speed (−3.6%) (**Fig. 3c**). Together, these findings indicate reduced vascular occupancy and structural narrowing on the stroke side, consistent with impaired microvascular perfusion despite preserved average flow speed at this acute stage after ischemic injury [33].

In the LPS-induced neuroinflammation model, co-registered ULM and PA imaging acquired 24 h after LPS administration showed that BBB leakage was scattered and bilaterally distributed across the cortex (**Fig. 3d**). Flow-speed–encoded ULM images from representative cortical ROIs likewise showed lower velocities after LPS treatment within individual brains (**Fig. 3e**). Whole-brain analysis across animals further demonstrated reduced blood volume and mean blood-flow speed after LPS treatment (**Fig. 3f**, paired t-test, n = 3 animals, p < 0.05), in agreement with prior Doppler ultrasound and MRI studies reporting reduced cerebral perfusion after systemic LPS challenge [34].

Together, our findings indicate that transient ischemia and systemic inflammation produce distinct microvascular phenotypes despite both causing BBB disruption. Stroke is associated with microvascular remodeling and region-dependent hemodynamic responses, whereas LPS causes diffuse BBB leakage accompanied by global suppression of microvascular perfusion. More broadly, these results show that 3D-PAULM can jointly map BBB permeability, vascular structure, and hemodynamics deep in brain.

### Reversibly switchable PAT (RS-PAT) enables background-suppressed tumor detection

We next evaluated the molecular imaging capability of 3D-PAULM in a tumor model using a genetically encoded photochromic probe (**Fig. 4a**). In conventional PA imaging, tumor signals are often obscured by surrounding blood vessels because hemoglobin dominates background absorption. To overcome this limitation, so-called photosensory core modules (PCMs) of bacterial phytochrome photoreceptors (BphPs) are used in the reversibly switchable PAT (RS-PAT) for background-suppressed tumor detection (**Fig. 4a**). PCM is a minimal part of BphP that preserves its photochromic behavior [35–37]. This temporal-modulation strategy isolates reversibly photoswitching probe signals from strong, non-switching background absorbers. For this, 4T1 breast cancer cells transfected with DrBphP-PCM were implanted subcutaneously into the left mammary gland of BALB/c mice to establish a BphP-expressing tumor model (**Fig. 4b**). Because tumor cores are typically poorly vascularized and surrounded by dense peripheral vessels, conventional hemoglobin-weighted PA images provide limited contrast for identifying the solid tumor core. To isolate the DrBphP-PCM signals, we implemented a photoswitching sequence on the 3D-PAULM system (**Fig. 4c**). Each 12-s cycle consisted of 6 s of 635-nm continuous-wave illumination to drive DrBphP-PCM into the ON state, followed by 6 s of pulsed 750-nm illumination, which both switched the probe toward the OFF state and served as the PA excitation light source. This sequence generated time-series data spanning alternating ON and OFF states.

**Fig. 4.**
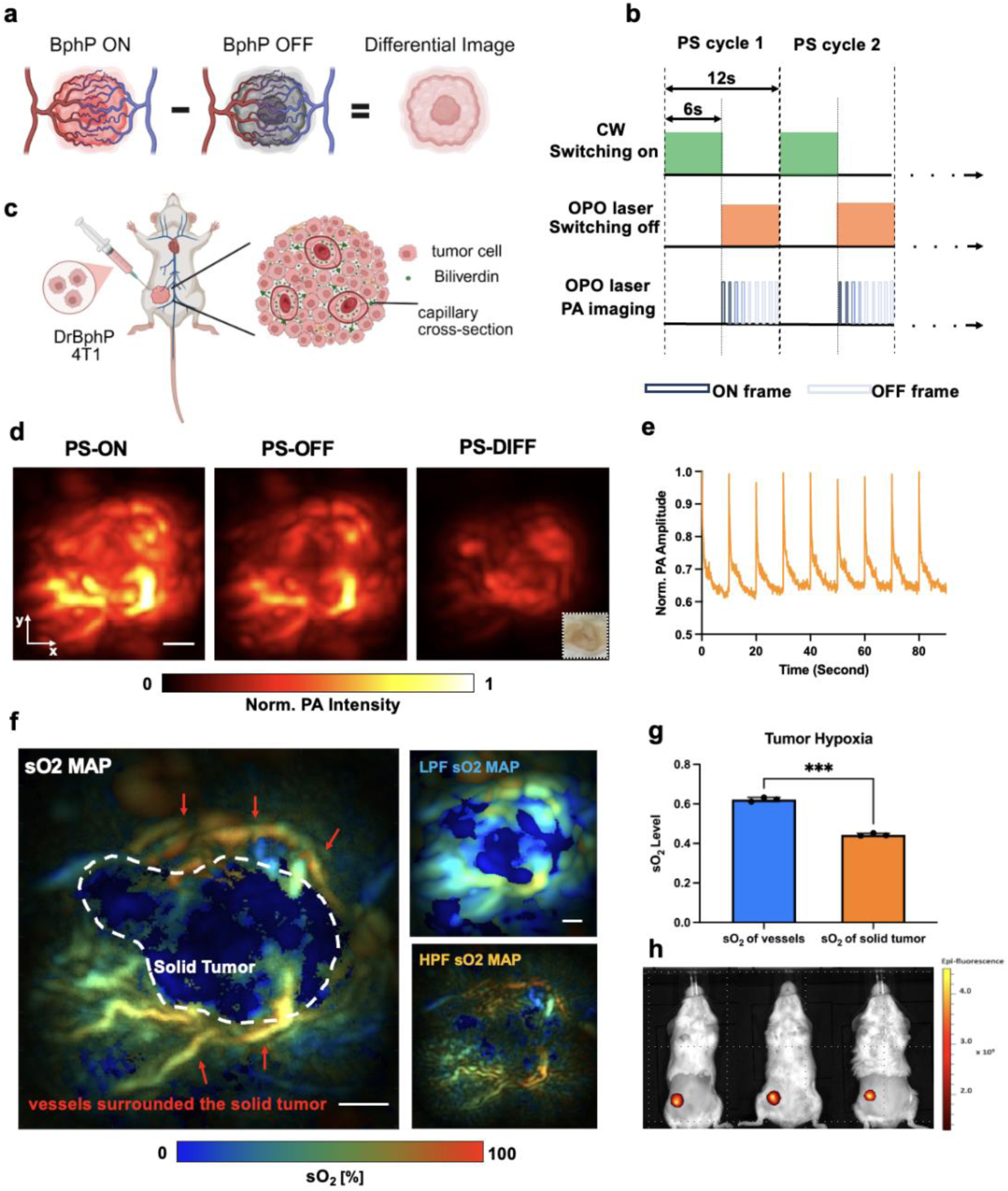
Functional and molecular photoacoustic imaging of BphP-expressing tumors using the 3D-PAULM system. **(a)** Principle of photoswitching (PS) PA imaging of a DrBphP-PCM-expressing tumor. **(b)** Imaging sequence of photoswitching acquisitions. A full PS cycle lasts 12 s, including 6 s switching ON and 6 s switching OFF periods. **(c)** DrBphP-PCM-transfected 4T1 breast tumor on the right mammary gland of a BALB/c female mouse. **(d)** *In vivo* photoswitching PA imaging. Representative PA images of the 4T1 tumor in PS-ON, PS-OFF, and differential (PS-DIFF) states. Scale bar, 2 mm. **(e)** Dynamic changes in photoswitching signals over nine photoswitching cycles. **(f)** Oxygen saturation (sO₂) mapping of the 4T1 tumor, showing marked hypoxia within the solid tumor core (white dashed line) compared with the surrounding vasculature (red arrows). Low-pass-filtered (LPF) maps emphasize bulk tumor oxygenation, while high-pass-filtered (HPF) maps highlight fine vascular structures. Scale bar, 2 mm. **(g)** Quantitative analysis of tumor hypoxia. Averaged sO₂ levels across three mice show significantly lower oxygenation in the tumor core than in the surrounding vessels (p = 0.0003). Data are presented as mean values ± SD (n = 3; independent transfection experiments). A paired t-test was used. **(h)** In vivo imaging of tumor-implanted mice using an IVIS Spectrum instrument. Excitation wavelength: 675 nm; emission wavelength: 703 nm.

As shown in **Fig. 4d**, both tumor tissue and vessels contributed to the PS-ON frames, whereas PS-OFF frames largely reflected the static hemoglobin background. Subtracting the OFF image from the ON image produced a differential (PS-DIFF) map in which non-switching background signals were strongly suppressed while photoswitchable-probe signals were retained. Several distinct foci then became visible in the PS-DIFF image, corresponding to tumor nodules. These RS-PAT image features closely matched the tumor nodules seen in the *ex vivo* photograph (**Fig. 4d**, inset). Time-course analysis across multiple photoswitching cycles showed consistent, reversible PA signal modulation (**Fig. 4e**), indicating stable probe switching during repeated acquisition. Together, these results demonstrate that DrBphP-PCM-based RS-PAT effectively suppresses hemoglobin-dominated background, thereby improving molecular contrast and enhancing detection of tumor-associated signals in deep tissue.

### Multispectral PA imaging reveals tumor hypoxia

Tumor hypoxia, a hallmark of aggressive solid tumors, reflects oxygen deficiency within the tumor core caused by abnormal and insufficient vasculature [38, 39]. We therefore used fluence-compensated dual-wavelength PA imaging (750 and 870 nm) to spectrally unmix hemoglobin species and quantify sO₂ in the DrBphP-PCM-expressing 4T1 tumor model. The reconstructed sO₂ maps, masked with PA images (**Fig. 4f**), revealed a pronounced hypoxic tumor core surrounded by relatively well-oxygenated peripheral vasculature.

Small, well-defined vessels preferentially contribute to high-frequency PA components, whereas the bulk, more homogeneous tumor contributes mainly to low-frequency components [40]. We therefore decomposed the PA data into high- and low-frequency bands. High-pass filtering (HPF, >4 MHz) emphasized fine vascular structures near the tumor surface, as reflected in the corresponding HPF sO₂ maps, whereas low-pass filtering (LPF, <2 MHz) enhanced signals from the deeper solid tumor mass in the LPF sO₂ maps. These observations support the use of low-frequency PA components for robust sO₂ quantification within the solid tumor core in this study.

Fluorescence imaging with IVIS confirmed the presence and location of DrBphP-PCM-expressing tumors on the ventral and left sides in all three mice (**Fig. 4h**). Statistical analysis of sO₂ in ROIs placed over tumor-surrounding vessels versus solid tumor cores showed significantly lower sO₂ in the cores (**Fig. 4g**; n = 3 animals, p < 0.001), further validating the presence of marked intratumoral hypoxia.

### ULM characterization of tumor vascular architecture and hemodynamics

Beyond sO₂ mapping, 3D-PAULM also enabled noninvasive visualization of vascular remodeling and hemodynamics in the DrBphP-PCM-4T1 tumor model (**Fig. 5**). Volumetric ULM of the bilateral lower abdominal mammary region, acquired with 2,500 frames per position to balance spatial coverage and total imaging time, revealed a striking vascular asymmetry: the mammary gland bearing the DrBphP-PCM-4T1 tumor exhibited markedly greater vessel density, tortuosity, and branching complexity than the contralateral side (**Fig. 5a**). Co-registered ULM and RS-PAT images further confirmed that the solid tumor was embedded within this angiogenic vascular network and enabled precise colocalization of molecular and vascular features.

**Fig. 5.**
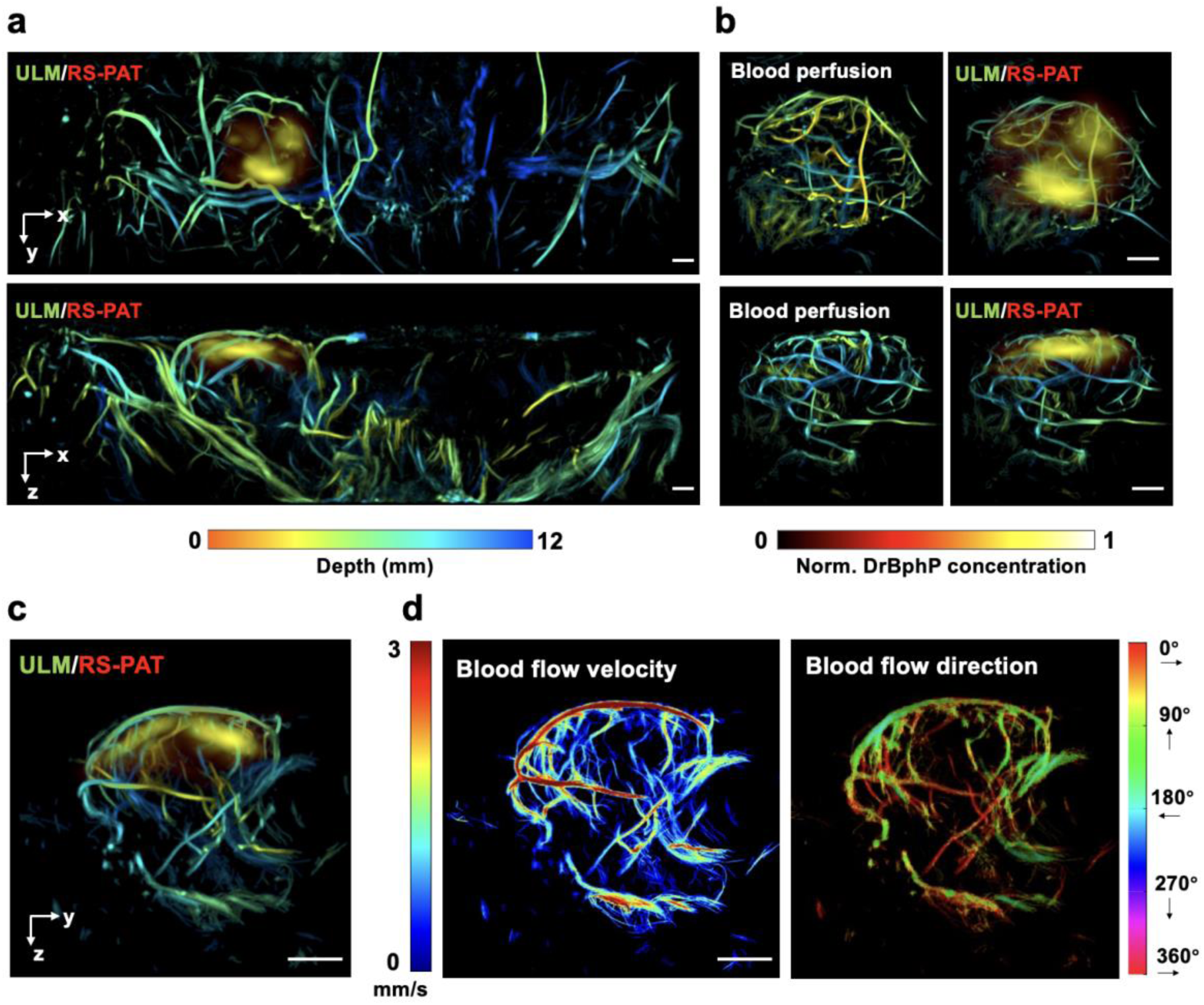
3D-PAULM imaging of the BphP-expressing 4T1 tumor. **(a)** Overlay of the ULM image showing vessels in the bilateral lower abdominal mammary region of a mouse bearing the DrBphP-PCM-4T1 tumor in the right mammary gland and the PA image showing the isolated DrBphP-PCM signal from the solid tumor. Maximum-intensity projections are displayed in transverse (top) and coronal (bottom) planes. **(b)** ULM blood-perfusion images of the tumor (left) and the corresponding ULM/RS-PAT overlays (right) for two orthogonal views. Compared with (a), the increased number of ULM frames yields richer microvascular detail in and around the tumor. **(c)** Overlay of the ULM image and RS-PAT of the tumor in the sagittal plane. **(d)** ULM maps of blood-flow velocity and direction in the tumor in the transverse plane. Scale bars, 2 mm.

To better resolve tumor-associated microvasculature, we then performed a tumor-focused ULM acquisition with 5,000 frames per position. The resulting blood-perfusion maps, together with the corresponding ULM/RS-PAT overlays, provided richer 3D vascular detail in and around the tumor (**Fig. 5b–c**).

Finally, ULM-derived blood-flow velocities and directions in the transverse plane (**Fig. 5d**) revealed heterogeneous flow behavior, including a broad range of microvessel flow speeds, slow-flow vessels interspersed among faster feeding vessels, velocity variations along individual vessel segments, and locally varying flow directions in neighboring branches. These observations are consistent with prior intravital imaging and modeling studies showing that tumor vasculature is often dilated, tortuous, and poorly hierarchized, leading to heterogeneous perfusion and flow dynamics [41].

Together, these results show that 3D-PAULM can integrate deep-tissue molecular contrast, oxygenation, and super-resolved vascular hemodynamics within a single experimental framework.

## Discussion

Here, we extend our previously reported 3D-PAULM platform from system development to biologically motivated applications, showing that integrated photoacoustic tomography and ultrasound localization microscopy can jointly interrogate deep-tissue molecular contrast and super-resolved hemodynamics in a noninvasive, co-registered manner. Building on the original hardware, we implemented faster data acquisition and streaming strategies to support high-throughput volumetric imaging. Combined with interleaved PA/US imaging at each scan position, these refinements improve spatiotemporal consistency across modalities, reduce motion artifacts, and shorten total image acquisition time. Comprehensive multiparametric imaging can be completed in under one minute per scan position, allowing whole-brain coverage within five minutes, which is important for application-driven *in vivo* studies requiring coordinated molecular, functional, and hemodynamic measurements.

Using this framework, we first demonstrated that multispectral PA imaging with ICG enables volumetric assessment of BBB disruption through intact skin and skull. Unlike fluorescence imaging, which is often biased toward superficial tissue, 3D-PAULM provides depth-resolved whole-brain maps of dye extravasation together with quantitative measures of leakage burden and spatial distribution. In the two brain models examined here, the volumetric imaging readouts distinguished focal, hemispherical asymmetric leakage after transient ischemia from diffuse, bilateral leakage after systemic LPS challenge. This ability to covert dye accumulation into three-dimensional leakage metrics is therefore a practical advantage of the platform for preclinical studies that require spatially resolved assessment of BBB opening or permeability changes over time.

By co-registering ULM with ICG-based PA images, 3D-PAULM places BBB permeability within its microvascular context, enabling barrier dysfunction to be interpreted alongside vascular structure and flow within the same tissue volume. The present findings further suggest that BBB disruption does not reflect a single vascular phenotype, but rather distinct modes of microvascular alteration. In transient ischemia, the shift toward smaller apparent vessel calibers, together with decoupling between vascular occupancy and flow metrics, suggests early microvascular reorganization rather than uniform perfusion loss. This pattern is consistent with a heterogeneous response involving local caliber narrowing, flow redistribution, and partial network-level compensation [32, 42]. By contrast, LPS-induced inflammation produced a more global impairment, characterized by concurrent reductions in blood volume and flow speed and indicative of widespread suppression of microvascular perfusion.

Notably, BBB disruption in both models occurred without overt infarction or structural tissue damage detectable by conventional imaging, indicating that barrier breakdown can arise from vascular dysfunction alone rather than requiring gross anatomical injury. This observation underscores that permeability changes, when considered in isolation, may not fully capture the underlying vascular state. Integrating permeability with microvascular structure and flow therefore provides a more specific framework for interpreting neurovascular pathology, with potential implications for understanding neurovascular coupling and optimizing therapeutic delivery [43, 44]. More broadly, the co-registered combination of molecular contrast and super-resolution hemodynamics enables *in vivo* interpretation of BBB leakage within its vascular context. Although the present study is limited by sample size and within-animal comparisons and estimates of vessel caliber may be influenced by ULM localization and tracking characteristics, the results support the potential of integrated PA/ULM imaging to resolve microvascular mechanisms of BBB dysfunction.

We next demonstrated a second key capability of 3D-PAULM by using photoswitchable probes for background-suppressed molecular imaging through differential photoacoustic detection. This approach complements the BBB studies by addressing the challenge of limited molecular sensitivity in blood-rich tissue. Temporal modulation of probe absorption allows RS-PAT to suppress non-switching background signals such as hemoglobin and recover weak molecular signals that are otherwise obscured in conventional PA imaging. The strategy is particularly well suited to genetically encoded probes, which enable cell-specific expression and support longitudinal *in vivo* studies without repeated administration of exogenous agents. More broadly, these results highlight temporally modulated contrast as a general strategy for improving molecular sensitivity in optoacoustic imaging.

We further used multispectral PA imaging to estimate sO₂ and provide functional context for tumor characterization. In the BphP-expressing 4T1 model, we observed a hypoxic tumor core surrounded by relatively well-oxygenated peripheral vasculature. This spatial mismatch between high peripheral vascular density and a hypoxic interior is consistent with the well-established supply-demand mismatch of aggressive solid tumors [45, 46].

Methodologically, the frequency-dependent analysis highlights a practical trade-off in deep-tissue PA imaging: low-frequency components are dominated by the bulk tumor, whereas higher-frequency components emphasize fine vascular features. Prioritizing low-frequency components for sO₂ estimation can therefore reduce the influence of superficial vascular heterogeneity and improve the robustness of deep-tumor oxygenation mapping, albeit at the cost of spatial resolution.

Although ULM confirmed an extensive angiogenic network around the tumor, the lower sO₂ within the tumor core suggests that vascular presence does not necessarily translate into effective oxygen delivery. By integrating molecular BphP-derived probe with functional oxygenation maps, 3D-PAULM enables interrogation of the metabolic–structural interface within the tumor microenvironment, which is important for studying tumor progression and responses to hypoxia-targeted therapies.

Despite these advances, several limitations remain. The 4-MHz hemispherical array, while enabling deep penetration and transcranial imaging, imposes finite spatial resolution and angle-dependent sensitivity on PA measurements [47]. In addition, quantitative interpretation of PA-derived molecular and functional signals remains affected by wavelength-dependent optical attenuation and fluence heterogeneity in deep tissue. Further improvements in fluence compensation and spectral unmixing will therefore be important for enhancing quantitative accuracy [48, 49].

ULM also requires microbubble contrast and relatively long acquisition windows. Although the higher volumetric frame rate and optimized data-streaming strategy reduce per-position acquisition time, motion artifacts and microbubble destruction remain potential challenges, especially in highly dynamic organs such as the lung or heart. Advances in motion correction, adaptive microbubble administration, and acquisition optimization will be important for extending 3D-PAULM to more motion-prone settings.

In summary, 3D-PAULM provides an integrated framework for multimodal deep-tissue imaging by combining molecular contrast, vascular structure, and hemodynamics within a single, intrinsically co-registered acquisition. By unifying multispectral photoacoustic imaging with super-resolution ultrasound localization microscopy, the platform enables simultaneous three-dimensional characterization of complementary biological parameters that are typically measured separately. Relative to our previous implementation, the present system offers improved imaging efficiency and spatiotemporal consistency across modalities, facilitating application-driven *in vivo* studies. Rather than being limited to a single biological question, 3D-PAULM offers a flexible strategy for linking molecular, functional, and microvascular information in complex biological systems. Together, these advances position it as a versatile tool for preclinical imaging and a foundation for future quantitative and potentially translational development in integrated functional and molecular imaging.

## Methods

### 3D-PAULM system and dual-mode acquisition

As shown in **Fig. 1a**, the 3D-PAULM system comprises a programmable 256-channel data acquisition system (Vantage 256, Verasonics, Inc.), a Q-switch laser equipped with an optical parametric oscillator (Innolas SpitLight I200 OPO, Amplitude Laser), and a customized 2D hemispherical ultrasound transducer array (Imasonics, France). Light was delivered via a fiber bundle positioned at the center of the array. Laser firing, ultrasound detection, and data acquisition were synchronized using a custom FPGA-based control module. During imaging, the transducer array faced upward within a temperature-controlled water bath. Imaging targets were mounted on an optically and acoustically transparent holder and raster scanned using a two-axis motorized translation stage.

A “ping-pong” data-saving scheme was implemented to decouple data acquisition from disk writing [50]. In this approach, two memory buffers were alternately used, allowing data acquisition to resume immediately after buffer transfer while data writing proceeded asynchronously. Interleaved PA and US acquisition was performed at each scanning position to ensure intrinsic spatial and temporal co-registration between modalities.

To accumulate sufficient frames for ULM while minimizing spatial misalignment between PA and US data, multiple cycles of alternating US and PA acquisitions were performed. US imaging employed synthetic aperture with 31-element transmissions at a center frequency of 4 MHz. For each transmission, backscattered echoes from all 256 elements were sampled at 15.625 MHz, and RF data were transferred in batches, resulting in a volumetric US frame rate of 403 Hz.

PA imaging was performed with multi-wavelength excitation at a pulse repetition frequency of 30 Hz. PA signals were sampled at 20.83 MHz and averaged over multiple laser shots to improve signal-to-noise ratio. The system supports acquisition of multispectral PACT, US B-mode, microbubble-enhanced power Doppler (PWD), and ULM data within a single scanning.

### Multi-spectral PA imaging and spectral unmixing

Multi-spectral PA imaging was performed to quantify hemoglobin oxygen saturation (sO₂) and relative indocyanine green (ICG) concentration. The OPO laser switched wavelengths every 30 firings. A fraction of the laser output was directed by a beam splitter to an energy meter, and the recorded energy values were used for pulse-by-pulse energy normalization. At each scan position, 30 laser pulses per wavelength were averaged to improve signal-to-noise ratio. Volumetric PA images were reconstructed using a three-dimensional delay-and-sum (DAS) algorithm with an isotropic voxel size of 60 μm, and volumes from all scan positions were stitched to generate the final 3D dataset.

Two excitation wavelengths (750 and 870 nm) were used for sO₂ estimation. For ICG mapping, three wavelengths (730, 800, and 870 nm) were selected to maximize spectral separability between absorbers while maintaining small inter-wavelength spacing. Molecular concentrations were estimated by fitting energy-normalized PA signals to reference absorption spectra using a linear least-squares method. sO₂ was calculated as the ratio of oxygenated hemoglobin to total hemoglobin. For ICG, reference spectra of oxyhemoglobin, deoxyhemoglobin, and ICG were used, with the ICG spectrum taken from plasma measurements reported by Landsman [27].

### Differential PA imaging with genetically encoded photochromic probes

BphPs incorporate endogenous biliverdin to form a chromophore that enables reversible photoswitching between two spectrally distinct absorbing states [21, 22]. This property allows background suppression in PA imaging by subtracting signals acquired in different absorption states.

A photoswitching PA imaging sequence was implemented using a DrBphP-PCM BphP. A 635 nm continuous-wave (CW) diode laser and the OPO laser were combined using a dichroic mirror and coupled into the fiber bundle. The 635 nm CW illumination was used to switch the probe to the ON state, whereas pulsed 750 nm illumination served both as PA excitation and to drive the probe toward the OFF state. Each photoswitching cycle lasted 12 s, consisting of 6 s of 635 nm illumination followed by 6 s of pulsed 750 nm illumination. Reconstructed PA volumes contained interleaved ON and OFF states. Differential images were obtained by subtracting the time-averaged OFF stack from the corresponding ON stack on a voxel-by-voxel basis. RF data were deconvolved with the measured electrical impulse response of the transducer array and low-pass filtered with a cutoff frequency of approximately 2 MHz. Multiple photoswitching cycles were averaged to improve signal stability.

### Microbubble-enhanced power doppler (PWD) and ULM

To visualize deep-brain and tumor hemodynamics at both conventional and super-resolution scales, we performed microbubble-enhanced 3D power Doppler (PWD) and ULM. Lipid-shelled microbubbles (Definity, Lantheus Medical Imaging) were activated by mechanical agitation (Vialmix activator, Lantheus Medical Imaging) according to the manufacturer’s instructions and diluted in sterile saline immediately before use to a final concentration of 7 × 10⁸ microbubbles mL⁻¹. For each imaging session, approximately 100 µL of the diluted microbubble suspension was administered as a bolus via the tail vein right before ULM acquisition (mouse body weight ∼25 g).

Volumetric US data for PWD/ULM were acquired using a synthetic-aperture transmission scheme mentioned above. Each transmitting element emitted a two-cycle burst at the center frequency. For deep-brain imaging, the driving voltage was set to 30 volts to compensate for the strong attenuation introduced by the skull, thereby improving the transcranial SNR. For superficial tumor imaging, the driving voltage was reduced to 15 volts instead, which was sufficient to maintain image quality in soft tissue while minimizing microbubble destruction near the surface. Backscattered echoes were recorded using all 256 elements for each transmission. For both brain and tumor PWD/ULM, over 2500 consecutive 3D US frames were acquired per scanning position. To reduce motion artifacts associated with respiration, lateral maximum-intensity-projection (MIP) images were cross-correlated frame by frame with an averaged reference MIP, and frames with correlation below 0.99 were discarded.

PWD images were generated following the same processing pipeline as in our previous 3D-PAULM work. Briefly, a spatial–temporal singular value decomposition (SVD) filter was applied to the DAS-reconstructed in-phase, quadrature (IQ) data to suppress tissue clutter. Higher-ranked singular values corresponding to relatively slow-varying and uniform structures (e.g., tissues) were set to zeros during reconstructions, leaving only singular values that match weak, fast-varying signals (e.g., flowing MBs, blood cells, etc.). The power of the filtered IQ volumes was then integrated over time to form the final PWD results. To further enhance separation of MB signals from noise and residual clutter, directional filtering and temporal band-pass filtering (20–107 Hz) were applied before integration [51], and the resulting PWD images were used as vascular masks for subsequent ULM analysis.

For ULM, centroids of SVD-filtered MB signals were localized at a subwavelength precision in each frame using a radial-symmetry-based localization algorithm [52]. Localizations from all frames were accumulated to generate super-resolved 3D vascular density maps. To estimate flow velocity and direction, individual MBs were tracked across successive frames using a Kalman filter-based 3D particle-tracking algorithm [53]. Localizations in consecutive frames were linked if they fell within a maximum search radius of 10 voxels (corresponding to a maximum velocity on the order of 100 mm/s) and a maximum gap of one frame. The displacement of each track over time yielded 3D velocity vectors, which were then used to compute flow speeds and flow directions.

Again, owing to the interleaving data acquisition scheme, misalignments between PA and US data remain minimal, which enables joint analysis of blood volume, flow velocity, and molecular contrast in both deep brain and tumor experiments.

### ULM image analysis

To quantify microvascular properties, flow speed, vessel diameter, and blood volume were further derived from the ULM data within manually defined regions of interest (ROIs). Flow speed was obtained from microbubble tracking, where the velocity of each tracked microbubble was calculated from frame-to-frame displacement, and flow-speed distributions were generated by aggregating all trajectories within each ROI.

Vessel diameter was estimated from the super-resolved ULM vascular density maps. Binary vessel masks were first generated from accumulated microbubble localizations using intensity thresholding. Vessel centerlines were then extracted via skeletonization, and local vessel radius was computed using a distance transform from the centerline to the vessel boundary. Vessel diameter was defined as twice the local radius, and diameter distributions were obtained by pooling all vessel segments within each ROI.

Blood volume was quantified as vessel volume fraction within each ROI, defined as the ratio of vessel voxels to the total number of voxels in the ROI based on the binary vessel mask.

### Blood brain barrier leakage mouse model

Two mouse models were used: (1) transient focal cerebral ischemia and (2) systemic lipopolysaccharide (LPS)–induced neuroinflammation. Female adult BALB/c mice (8–12 weeks) were used. All procedures were approved by the Duke University Institutional Animal Care and Use Committee (IACUC) and complied with institutional animal care guidelines.

Baseline imaging was performed prior to any surgical or inflammatory intervention. For baseline imaging, animals were anesthetized (1.5–2% isoflurane in oxygen) and intravenously injected with indocyanine green (ICG) to achieve a final plasma concentration of approximately 13.5 μM. PA/ULM imaging was performed 20 min after injection. Twenty-four hours after induction of transient focal ischemia or systemic LPS treatment, mice received a second intravenous tail-vein injection of ICG at the same dose (13.5 μM final plasma concentration). PA/ULM imaging was performed 20 min after injection. For the transient focal ischemia cohort, mice were subsequently transferred to a fluorescence imaging system immediately after 3D-PAULM imaging for additional validation.

In the transient focal ischemia model, unilateral occlusion of the right carotid circulation was used to induce focal BBB leakage. Mice were anesthetized and placed on a temperature-controlled heating pad to maintain core body temperature. After a midline cervical incision, the right common carotid artery was isolated from the vagus nerve and surrounding connective tissue, and a microvascular clip was applied to interrupt blood flow to the ipsilateral (right) hemisphere. During occlusion, respiratory rate and pedal reflexes were monitored, and anesthetic depth was adjusted as needed. After 30 min of occlusion, the clip was removed to allow reperfusion; hemostasis was confirmed, and the incision was closed with absorbable sutures. Mice were returned to a warmed recovery cage and monitored until fully ambulatory. Animals showing severe distress or failing to recover were excluded according to predefined humane endpoints.

In the systemic LPS-induced neuroinflammation model, mice were anesthetized briefly for handling, weighed, and injected intraperitoneally with LPS (Escherichia coli O111:B4, Sigma) dissolved in sterile water at a dose of 5 mg/kg. After injection, animals were returned to their cages with free access to food and water and monitored for systemic sickness behavior. Imaging was performed 24 h after LPS administration following the same ICG injection and 3D-PAULM protocol described above.

### Photoswitchable tumor mouse model

A subcutaneous breast tumor model expressing the DrBphP-PCM was established for in vivo molecular imaging. Murine 4T1 breast cancer cells were engineered to stably express DrBphP-PCM through lentiviral transduction. Cells were infected with a lentiviral vector encoding the photosensory core module of DrBphP-PCM linked to EGFP via an internal ribosome entry site. Forty-eight hours after infection, EGFP-positive cells were enriched by fluorescence-activated cell sorting and expanded to generate a stable DrBphP-PCM-4T1 cell line. Cells were maintained in DMEM (Thermo Fisher Scientific) supplemented with 10% fetal bovine serum and 1% penicillin–streptomycin at 37 °C in a humidified 5% CO₂ incubator and passaged before reaching confluence. Cells between passages 3 and 10 were used for *in vivo* experiments.

For tumor implantation, female BALB/c mice (6–8 weeks) were anesthetized with 1.5–2.0% isoflurane in oxygen and placed in the supine position. Fur over the fourth mammary fat pad was shaved and disinfected with povidone–iodine followed by 70% ethanol. A suspension of DrBphP-PCM-4T1 cells in sterile PBS (1×10⁷ cells mL⁻¹) was prepared on ice, and 0.1 mL (1×10⁶ cells) was injected into the right fourth mammary fat pad using a 27-gauge needle. Gentle pressure was applied at the injection site for several seconds. Mice were monitored daily, and tumors were imaged when they reached approximately 8 mm in diameter (typically 10–14 days post-implantation). At the end of imaging, animals were euthanized according to IACUC-approved protocols, and tumor samples were collected for ex vivo fluorescence validation.

## Data availability

All data supporting the findings of this study are available within the paper. Source Data is provided with request to the corresponding authors.

## Code Availability

The image processing codes used in this study are available at Duke Photoacoustic Imaging Lab’s GitLab page: https://gitlab.oit.duke.edu/pilab/3D-PAULM

## Acknowledgments

This work was partially sponsored by the United States National Institutes of Health (NIH) grants R01EB037095, R01ES036951, RF1 NS115581, R01 NS111039, R01 EB028143, R01 DK139109, R01 DK052985, R01 MH135932, R35 GM122567; The United States National Science Foundation (NSF) CAREER award 2144788; American Heart Association Collaborative Science Award (25CSA1417550); Duke Gilhuly Acceleration Grant; Duke University Pratt Beyond the Horizon Grant; Chan Zuckerberg Initiative Grant (2024-349531); Duke University DST Spark Seed Grant; Duke Coulter Translational Grant; North Carolina Biotechnology Center Triangle Research Grant (2024-TRG-0041); and Einstein 2030 Seed Fund.

## Author contributions

Y.X. performed the 3D-PAULM imaging and data analysis and assisted with the 3D-PAULM imaging system construction. H.S., X.Y. and W. Y performed animal surgeries for all the 3D-PAULM experiments. R.Y. and N.W. contributed to the image reconstruction algorithm for 3D-PAULM. P.S. developed the ULM algorithm. X.C. and J.C. assisted with the 4T1 cell culture. J.L. and J.L. assisted with animal breeding and fluorescence imaging. V.V.V. designed the genetically encoded photoswitching probe and related experiments. J.Y. designed the 3D-PAULM imaging experiments. J.Y. conceived the project. J.Y. and V.V.V. supervised the project. Y.X. wrote the manuscript. R.Y., and J.Y. reviewed and revised the manuscript.

